# Cancer3D 2.0: interactive analysis of 3D patterns of cancer mutations in cancer subsets

**DOI:** 10.1101/420521

**Authors:** Mayya Sedova, Mallika Iyer, Zhanwen Li, Lukasz Jaroszewski, Kai W. Post, Thomas Hrabe, Eduard Porta-Pardo, Adam Godzik

## Abstract

Our knowledge of cancer genomics exploded in last several years, providing us with detailed knowledge of genetic alterations in almost all cancer types. Analysis of this data gave us new insights into molecular aspects of cancer, most important being the amazing diversity of molecular abnormalities in individual cancers. The most important question in cancer research today is how to classify this diversity to identify subtypes that are most relevant for treatment and outcome prediction for individual patients. The Cancer3D database at http://www.cancer3d.org gives an open and user-friendly way to analyze cancer missense mutations in the context of structures of proteins they are found in and in relation to patients’ clinical data. This approach allows users to find novel candidate driver regions for specific subgroups, that often cannot be found when similar analyses are done on the whole gene level and for large, diverse cohorts. Interactive interface allows user to visualize the distribution of mutations is subgroups defined by cancer type and stage, gender and age brackets, patient’s ethnicity, or vice versa find dominant cancer type, gender or age groups for specific three-dimensional mutation patterns.

## INTRODUCTION

Availability of large cancer genomics datasets, such as The Cancer Genome Atlas (TCGA) (1) and growing computational infrastructure at NCI (https://gdc.cancer.gov), UCSC’s Cancer Genomics Browser (2) or cBioPortal (3,4) allows individual researchers to analyze this data through public tools or download it for tailored analyses. The TCGA data was analyzed by the TCGA associated research groups that published series of seminal papers on many aspects of cancer (https://cancergenome.nih.gov/publications). Cancer3D database (5) developed in our group was one of the pioneers in integrating genomic data such as missense mutations with structural information on domain and three-dimensional structure of proteins harboring these mutations. Many servers added or expanded such capabilities since then and tools such as cBioPortal (3,4) allow for visualization and analysis of cancer mutations in 3D as one of the options in broader analysis and dedicated servers such as MuPIT (6), hotspots3D (7) or cosmic3D (8) focus specifically on understanding cancer mutations in their 3D context. However, most of the existing resources, following the path of most TCGA analyses, focus on large cohorts of patients with specific cancers or even groups of cancers such as gastrointestinal or gynecological cancers, seeking general observations that could be applied to large percentage of patients. On the other hand, many clinical and epidemiological analyses suggest existence of smaller or alternatively defined subgroups of patients. For instance, many cancers have different incidence levels, presentations and outcomes in males and females (9), in younger and older patients (10) or in patients with different ethnic background (11) and in most cases, the research into the molecular reasons for these differences is still in its infancy.

To help answer such questions, in a new, completely reworked version of Cancer3D database described here we integrate data from TCGA with protein 3D information and partial clinical data, to allow users to explore cancer mutations in their three-dimensional context for subgroups defined by clinical information. Cancer3D interface is gene-centric, asking users to provide a gene name to start the analysis. Similarly to the previous version of Cancer3D, every gene page provides information on functional modules within proteins: Pfam domains (12), set of PDB (13) coordinates mapped on a given gene, predicted intrinsically disordered regions, and, new in this version, interaction interfaces and structurally flexible regions of a given protein. The Cancer3D database does not only displays the mutated positions of a protein in their corresponding structures but allows users to interactively visualize individual positions using a system of pull down menus (Figure 1).

**Figure 1.**
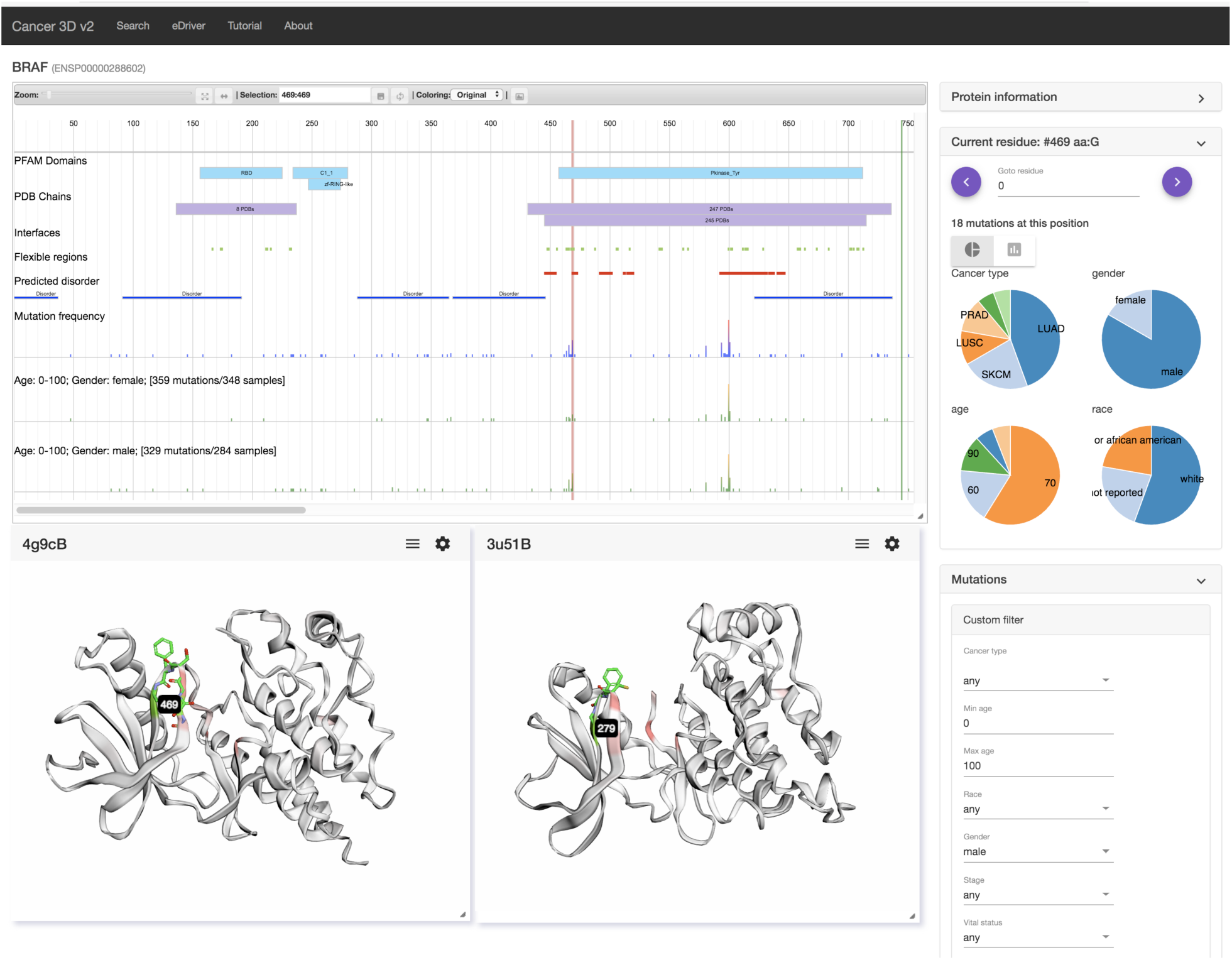
Sample output of Cancer3D database illustrating the example discussed in the text. Position #469 in the cluster of mutations on the glycine rich region is highlighted, panels showing gender, cancer type, age and ethnicity distributions are show on the left, two 3DMol windows illustrating the position of the 469 residue are shown in the lower part of the figure. The exact steps needed to achieve this output are outlined in the text. The on-line tutorial provides more detailed discussion of all the options in the database.

## MATERIAL AND METHODS

### Data sources

Mutation data used in Cancer3D database come from the Public MC3 MAF (13) distributed by The Cancer Genome Atlas analysis project (14) and sample information (age, gender, clinical state) comes from NCI Genomic Data Commons (https://gdc.cancer.gov). Variant Effect Predictor Tool was used to map mutations from genomic coordinates to all ENSEMBL proteins sequences (15). Cancer3D has a total of 1,457,702 mutations from 9079 samples representing 32 cancer types and mapped into 18,425 unique proteins.

Protein domains are assigned using Pfam HMM models as retrieved from ENSEMBL through its API, intrinsically disordered regions were predicted using Foldindex (16). Protein interaction interfaces were defined as in (17), in short for every set of PDB coordinates residues which had any heavy atom within 5A from a heavy atom from a different chain were marked as belonging to an interface. In the original paper we focused on protein-protein interfaces, now the same approach is extended to protein-ligand and protein-DNA interfaces. It’s important to note that interfaces are defined for each set of PDB coordinates separately and only then mapped on the gene, so every set of PDB coordinates can in principle have a different interface and the 2D protein overview page presents a sum of all interfaces, while in 3D visualization windows interfaces and other features are shown only for individual structures. Overall, 17,772 proteins contain one of the 6159 Pfam domains or a disordered region and 10,948 proteins have at least one of the structural features, such as an interaction interface or a flexibility region.

### Algorithms

We have previously developed an e-Driver algorithm (5) that allow the identification of candidate cancer drivers on the level individual domains. Later, we expanded this concept to apply it to protein-protein interfaces (17). In the current Cancer3D release we provide access to both domain and protein-protein interaction significance calculation and expand them to include intrinsically disordered regions, protein-ligand or protein-DNA interaction interfaces or regions with significant local flexibility. Collectively we refer to the interaction interfaces and flexible regions as “structural features” and we plan to expand the list of available features in the future releases of the Cancer3D database.

### Structure mapping

We have used BLAST (18) to match sequences of three-dimensional structures to all the genes in the database. The full PDB database was queried for each protein sequence in the Cancer3D database and e,value threshold of e-6 was used to match PDB structures to the proteins in the database. The BLAST output was used to map mutated positions onto the structures.

### Flexible regions mapping

Flexible regions for each protein mapped to a given gene are extracted from the PDBFlex database (19). In short, a sliding 10 residue window was used to calculate RMSD between a given protein and all proteins in the PDB with more than 95% sequence identity. Center residue of a window that has >1.5A RMSD in any comparison is tagged as “flexible”.

## USING THE DATABASE

### Use-case scenario

In the following paragraphs we will present an example of using the Cancer3D database for the analysis of mutations in the BRAF protein. This is a well-known oncogene with extensive annotations and analyses of many individual mutations (20).

### Input

The start page contains an input field to search for the user’s protein of interest. The user can input gene or protein name, Uniprot ID or ENSEMBL Gene / Protein ID. If the name is recognized by the database a 2D overview page will be displayed showing information for a “canonical” isoform of a given gene. A histogram of mutations and Pfam families, PDB structures and various structural features are mapped into the main protein sequence. Since we present the functionality of Cancer3D based on a use-case scenario using BRAF, our user would specify BRAF in the input menu. For the user interested in just browsing and getting familiar with the database we provide a link to a “random protein” that chooses a random protein from the database and displays its main annotation page. A number of analysis options are available either by utilizing information panels to the right of the main window or by moving a mouse over various icons on the page and selecting from the appearing drop-down menus.

### Main window

The main window shows the mutation histogram shown on a logarithmic scale, to allow frequently mutation positions to fit into the main window without resizing it. On the BRAF page, position V600, the most frequently mutated site, appears as the highest point in the histogram (See Figure 1). Above the histogram, blue boxes identify positions of Pfam domains, while purple ones denome positions of sets of PDB coordinates. Since different PDB files correspond to different constructs, they are grouped together when the constructs ends are within 5 residues of each other. For well-studied proteins only a number of PDB coordinates within given boundaries is show, if possible PDB IDs are shown in the box(se). Protein structural features are identified by colored boxes, blue for disordered regions, red for flexible regions and green for interaction interfaces.

Moving the mouse over the mutation histogram allows users to select specific position along the sequence of the protein. Detailed information about the mutation at a selected location would appear in the “Current residue” panel on the right, which may have to be open by clicking on the “>” sign. A number of mutations on a selected position is shown, together with histograms (shown as a standard histogram or as a pie chart) of cancer types, gender ratio, and race and age distribution of patients with mutation at this position. Instead of moving the mouse, a specific position can be selected by providing its position in the ENSEMBLE numbering scheme in the “Current residue” panel or by moving the current residue position using arrows in the top part of the current residue window.

For our use-case scenario for BRAF, the user would use the custom filters available in the “Mutations” panel on the right to select histograms for “female” and “male” available under the “gender” drop down menu. Two additional histograms appear in the main window. Other choices in the mutations window include selecting specific cancers, age groups, race, cancer stage or survival status of the patients. On the two new histograms it is possible to notice significant differences between the male and female histograms in several places (they may be easier to see using the zoom option and then sliding the view window along the length of the protein). For instance, there are no mutations in female patients in the region corresponding to the C1_1 domain and the group of mutations around residues 465-470 come predominantly from male patients. The latter group also corresponds to an interface (green box) in several proteins – their ID could be seen by moving the mouse over the green box in the main window. Selecting position 469 for a detailed analysis, we see that indeed 15 of the patients with mutations on this position are male, while only 3 are female. From other histograms in this panel we see that a significant number of patients were diagnosed with lung cancer (LUAD and LUSC) which is slightly more common in men and two cases were prostate cancer, but other cancers were melanoma (SKCM) and other non-gender specific cancer, so cancer distribution cannot explain the skewed male/female ration. Overall among the patients with mutations in the 465-470 regions, 22 were male and 6 were female.

### 3D windows

Using pop-up menus that become visible when moving a mouse over annotations in the main window, a user can select individual PDB files to detailed view. For instance, from the main purple box with 247 PDBs, we can select 4g9cB set of coordinates. This action opens a 3Dmol (21) window with an interactive 3D visualization of the kinase domain of BRAF, with initial coloring corresponding to the mutation frequency distribution. In our user scenario, a user can open a second window using the “interface” feature box. Chicken src kinase is a homolog of human BRAF and coordinate set 3u51B shows a complex of the scr kinase with an inhibitor, highlighting a possible function of this protein region. The 3DMol windows can we resized, the structures can be rotated and the CHIMERA {Pettersen, 2004 #23} script for further, more detailed analysis can be downloaded. The main window and the 3D view window(s) are integrated, selecting the position 469 on the histogram (or in the current residue window) highlight the corresponding residue in the 3D view and allow to study its structural context. The region 465-470 corresponds to the well conserved glycine-rich loop of the kinase domain with the GXGXXG motif, which is universally conserved in protein kinases and other proteins that bind mono- and dinucleotides (20). Mutations in this motif, especially the G469E mutation were extensively studied and associated mostly with lung cancer. Their strong gender bias was not, to the best of authors knowledge, discussed in literature.

## CONCLUSIONS AND FUTURE DEVELOPMENTS

We developed the Cancer3D database to allow users to analyze patterns of distribution of cancer mutations in the context of their three-dimensional structures integrated with information on clinical correlates of such patterns. Ethnicity, gender and age strongly correlate with specific course and outcomes of many kinds of cancer and a possibility of mutation patterns, and by extension, molecular mechanisms in tumors specific to these subgroups have not been deeply explored. We believe that the new Cancer3D database gives such option to the cancer research community.

## ACKNOWLEDGEMENTS

Authors want to acknowledge the TCGA consortium and NCI Data Commons for providing open source data that have been used in the Cancer3D database.

